# Determination of HLA-DR level in cytotoxic T lymphocytes: a new validated tool to predict breast cancer response to treatment

**DOI:** 10.1101/2021.04.29.441985

**Authors:** D.P. Saraiva, S. Azeredo-Lopes, A. Antunes, R. Salvador, P. Borralho, B. Assis, I. Pereira, Z. Seabra, I. Negreiros, A. Jacinto, S. Braga, M. G. Cabral

## Abstract

**Background:** Neoadjuvant chemotherapy (NACT) is the usual treatment for locally-advanced breast cancer (BC). However, more than half of the patients lack an effective response to this treatment. Thus, it’s urgent to find predictive biomarkers. Recently, we proposed the HLA-DR expression level in cytotoxic T lymphocytes (CTLs) as a robust biomarker to select, in advance, patients that will actually benefit from NACT.

**Patients and Methods:** A total of 202 BC patients, 102 of which submitted to NACT, were enrolled in this study. 61 biopsies and 41 blood samples collected pre-NACT and 100 non-NACT tumor samples were immunophenotyped by flow cytometry. Both NACT and non-NACT patients were followed up for 34 months. Blood-isolated immune cells were cultured with BC cell lines in a 3D system.

**Results:** Here we confirmed that HLA-DR level in CTLs is a highly sensitive and specific biomarker to predict BC response to NACT, reflected in circulation and independent of the patients’ age, BC subtype and other tumor-immunological features. Therefore, we developed a predictive probability model, based on the determination of HLA-DR level in tumor-infiltrating CTLs, that could be used to guide therapeutic decisions. Interestingly, this biomarker was also associated with progression-free survival, regardless the treatment. Contrary to HLA-DRnegative CTLs, HLA-DR+ CTLs were able to reduce the viability of tumor cells, in culture, in agreement with their higher expression of activation, proliferation and cytotoxicity-related molecules. Tissue-residency and memory markers were also increased in HLA-DR+ CTLs. These anti-tumor features of HLA-DR+ CTLs may justify the clinical observations.

**Conclusion:** HLA-DR level in CTLs is a validated and independent biomarker to predict response to NACT which allow the establishment of a clinical meaningful tool to select in advance patients that will truly benefit from this treatment. Intriguingly, it may be further used as a biomarker of BC patients’ general prognosis.

**Highlights:**

- HLA-DR level in cytotoxic T cells is an independent predictive factor of breast cancer response to NACT
- A predictive probability model based on this biomarker was developed as a new tool to improve treatment decisions
- HLA-DR level in cytotoxic T cells is also reflected systemically
- HLA-DR level in cytotoxic T cells is also a prognostic factor, associated with progression-free survival
- HLA-DR+ cytotoxic T cells exhibit several phenotypic and functional anti-tumor characteristics

## Introduction

Breast cancer (BC) is the most frequent type of cancer in women worldwide, accounting for up to 2 million new cases per year (1). Early-stage disease has a high survival rate, with 99% for estrogen receptor positive (ER+) tumors, 94% for human epidermal growth factor receptor 2 (HER2+) overexpressing tumors and 85% for triple negative breast cancer (TNBC)(2). Nevertheless, when the disease is in an advanced stage, the survival rate is lower, probably due to the lack of effective specific treatment options (2). Indeed, independently of the BC subtype, the treatment option for locally advanced BC is neoadjuvant chemotherapy (NACT). Although NACT is important in downstaging the tumor, allowing a breast-conserving surgery (3), approximately half of the patients do not respond to this treatment (4,5). Thus, it is essential to find biomarkers of response to NACT, as well as alternative therapeutic options for nonresponders.

NACT efficacy may depend on immune players present in the tumor microenvironment, namely tumor-infiltrating lymphocytes (TILs), and as such TILs have been studied as possible biomarkers to predict response to NACT. Actually, chemotherapy promotes immunogenic cell death (ICD), which involves the release of damage-associated molecular patterns (DAMPs) by dying cells, that might lead to the activation of particular TILs (namely cytotoxic T lymphocytes – CTLs) that, in turn, will allow the control of NACT-resistant tumor cells (6). Though accumulating evidence has been shown that the same ICD-resulting DAMPs could, in certain conditions, support cancer progression and resistance to treatment (7). Additionally, tumors have several mechanisms to escape immune surveillance and TILs-mediated killing (8). Hence, quantification of TILs is still not used routinely in the clinical scenario.

We have previously reported that CTLs expressing high levels of the activation marker HLA-DR, in fresh BC biopsies, were a highly sensitive and specific factor to predict response to NACT (9). We found that HLA-DR+ CTLs were mainly present inside the tumor microenvironment; produced high levels of cytotoxicity-related molecules as IFN-γ, Granzyme B and Perforin; were negatively correlated with immunosuppressive features of the tumor milieu and were reflected systemically (9). In order to confirm HLA-DR-expressing CTLs as a biomarker of response to NACT, following the REMARK (reporting recommendations for tumor marker prognostic studies) criteria (10), we conducted a validation study in an independent cohort of BC patients, tackling one of the clinical needs in BC management. Moreover, we took advantage of a 3D coculture platform of BC cell lines and patient-derived immune cells, developed to be used in the future as a drug screening platform (11), to reveal that HLA-DR+ CTLs, present in the blood of NACT-responders, are indeed cytotoxic towards tumor cells.

Thus, here we validate HLA-DR level in CTLs as a new, independent biomarker to predict BC response to NACT and additionally as a BC general prognosis factor. We further developed a predictive probability model, which we believe is a valuable tool to help clinicians to segregate NACT-responders from non-responders, in advance. Moreover, we unveil some phenotypic and functional characteristics of HLA-DR+ CTLs that explain, at least in part, the relevance of these cells in assisting NACT.

## Materials and Methods

### Patients’ samples

This prospective study was designed following the REMARK criteria (10). A total of 202 breast cancer (BC) patients were enrolled in this study (Fig.1). 61 fresh biopsies (from a previous pilot cohort and from this validation cohort) of BC patients selected for neoadjuvant chemotherapy (NACT) were collected in Transfix (Cytomark) to preserve cellular antigens. NACT was composed of 4 cycles of doxorubicin and cyclophosphamide every 3 weeks, followed by 12 weeks of paclitaxel. Trastuzumab was given every 3 weeks to HER2+ patients. A summary of the patients’ characteristics is presented in Table 1. The biopsies were mechanically dissociated with a BD Medicon (BD Biosciences), filtered through a 30 μm mesh (Sysmex), washed with PBS 1X and stained for flow cytometry (as described below). Biopsies and surgical specimens from non-NACT patients were similarly handled and were used, together with NACT samples, in the progression-free survival analysis.

**Fig.1.**
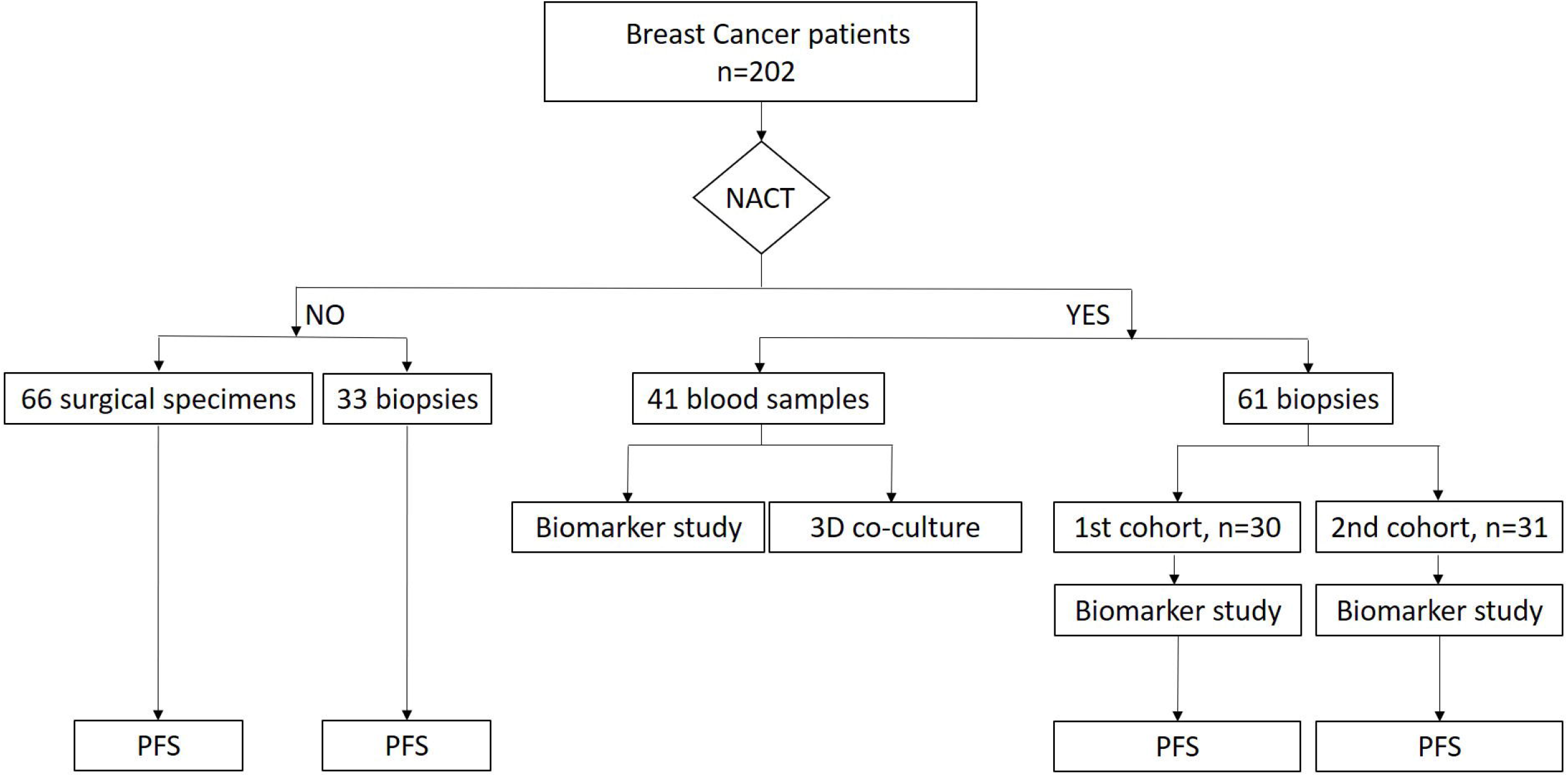
Flowchart of patients enrolled in this study. Breast Cancer patients (n=202) were divided according to the prescription of neoadjuvant chemotherapy (NACT). The biopsies of NACT patients were divided into two different cohorts for the predictive biomarker study. Blood samples were also used in this study and to isolate PBMCs for *in vitro* 3D co-culture assay. Progression-free survival (PFS) was performed for NACT and non-NACT patients.

**Table 1.**
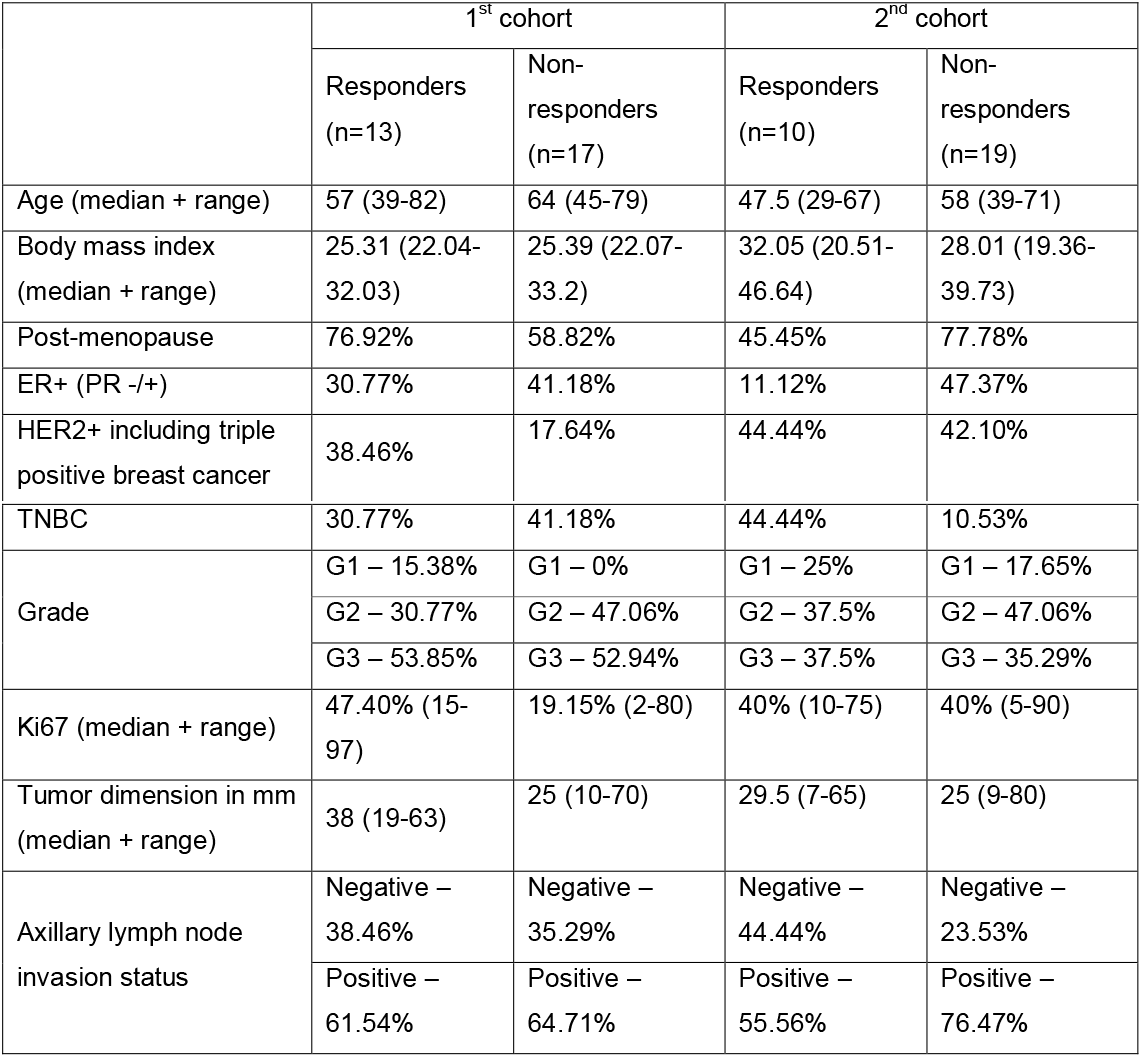
Characteristics of patients enrolled in this study (median values of age, body mass index and percentage of post-menopause patients). Clinical data, such as subtype of breast cancer, grade, median of Ki67 and of tumor dimension and node status are also described.

|  | 1<sup>st</sup> cohort |  | 2<sup>nd</sup> cohort |  |
| --- | --- | --- | --- | --- |
|  | Responders<br>(n=13) | Non-responders<br>(n=17) | Responders<br>(n=10) | Non-responders<br>(n=19) |
| Age (median + range) | 57 (39-82) | 64 (45-79) | 47.5 (29-67) | 58 (39-71) |
| Body mass index<br>(median + range) | 25.31 (22.04-32.03) | 25.39 (22.07-33.2) | 32.05 (20.51-46.64) | 28.01 (19.36-39.73) |
| Post-menopause | 76.92% | 58.82% | 45.45% | 77.78% |
| ER+ (PR -/+) | 30.77% | 41.18% | 11.12% | 47.37% |
| HER2+ including triple positive breast cancer | 38.46% | 17.64% | 44.44% | 42.10% |
| TNBC | 30.77% | 41.18% | 44.44% | 10.53% |
| Grade | G1 – 15.38% | G1 – 0% | G1 – 25% | G1 – 17.65% |
|  | G2 – 30.77% | G2 – 47.06% | G2 – 37.5% | G2 – 47.06% |
|  | G3 – 53.85% | G3 – 52.94% | G3 – 37.5% | G3 – 35.29% |
| Ki67 (median + range) | 47.40% (15-97) | 19.15% (2-80) | 40% (10-75) | 40% (5-90) |
| Tumor dimension in mm (median + range) | 38 (19-63) | 25 (10-70) | 29.5 (7-65) | 25 (9-80) |
| Axillary lymph node invasion status | Negative – 38.46% | Negative – 35.29% | Negative – 44.44% | Negative – 23.53% |
|  | Positive – 61.54% | Positive – 64.71% | Positive – 55.56% | Positive – 76.47% |

41 blood samples were collected from non-matched BC patients selected for NACT in Vacutainer tubes with EDTA (BD Biosciences). Whole blood staining for flow cytometry was performed as described below. Peripheral blood mononuclear cells (PBMCs) were isolated from whole blood by a Ficol gradient (Merck Millipore) and cryopreserved in 90% fetal bovine serum (FBS, Biowest) and 10% dimethyl sulfoxide (DMSO, Sigma Aldrich) to be used later in the 3D co-culture systems. Isolated PBMCs from 8 healthy donors’ buffy coats provided by Instituto Português do Sangue e da Transplantação were used specifically in the cytotoxicity assay.

Samples were gathered from Hospital CUF Descobertas, Hospital de Vila Franca de Xira, Hospital Prof. Doutor Fernando Fonseca and Hospital Santa Maria. For each patient, written informed consent and approval by the Ethical Committees of the hospitals and of the NOVA Medical School were obtained. The study complies with the Declaration of Helsinki.

### Flow cytometry

Cell suspension from the processed BC samples was stained with a cocktail of mouse anti-human surface antibodies for 15 min. The cells were then fixed and permeabilized with Fix/Perm kit (Invitrogen) followed by intracellular staining for 30 min. In case of whole blood, the staining protocol was similar, but it was followed by a step of red blood cells lysis with RBC lysis buffer (Biolegend). Data were acquired in BD FACS Canto II (BD Biosciences) and the results were analyzed using FlowJo software v10. The data are presented as a percentage of the populations in respect to the single cells’ gate. To analyze the expression levels of HLA-DR in cytotoxic T lymphocytes (CTLs), we considered the median fluorescent intensity of the positive population and normalized it to the negative population. The negative population was superimposed with the unstained control.

The monoclonal mouse anti-human antibodies used were: anti-CD45-PercP, anti-CD3-APC, anti-CD19-PE, anti-CD15-PE, anti-CD161-FITC, anti-CD4-FITC, anti-CD8-PE, anti-HLA-DR-APC, anti-CD127-PE-Cy7, anti-CD25-PE, anti-CD163-PE, anti-CD206-APC-Cy7, anti-PD-L1-APC, anti-CD11b-FITC, anti-IL-10-FITC, anti-CD69-PercP, anti-IFN-γ-APC-Cy7, anti-GranzymeB-FITC, anti-Ki67-PE, anti-CD45RO-PercP, anti-PD-1-FITC, anti-Tim3-APC-Cy7, anti-CD39-BV421 (Biolegend) and anti-CD103-PE-Cy7 (Invitrogen).

### Fluorescence activated cell sorting (FACS)

PBMCs from healthy donors were cultured overnight in RPMI-1640 (Gibco) supplemented with 10% FBS and 1% Penicillin/Streptomycin (GE Healthcare) and stimulated with 35 ng/mL of phorbol 12-myristate 13-acetate (PMA, Sigma Aldrich) and 1 μg/mL of ionomycin (Merck Millipore) at 37°C, 5% CO_2_. Cells were then stained with BD Horizon^™^ Fixable Viability Stain 450 (BD Biosciences) for 20 min in ice, followed by staining with the antibodies anti-CD45-PercP, anti-CD8-PE and anti-HLA-DR-APC. Cells were sorted into two populations: CD45+/CD8+/HLA-DR+ and CD45+/CD8+/HLA-DRnegative in a FACS Aria III (BD Biosciences) with an efficiency above 90%.

### Cytotoxicity assays in 3D co-cultures

The BC cell lines MCF-7 and MDA-MB-231 were cultured in DMEM (Biowest) supplemented with 10% FBS and 1% Penicillin/Streptomycin. 3D co-culture in agarose-coated plates of both cell lines with patient-derived PBMCs in a 1:1 ratio was performed as previously described (11). 4 days later, the spheroids were removed from the plate, dissociated by pipetting and stained with the Viability Dye to evaluate, by flow cytometry, the percentage of viable cancer cells in the co-culture.

Additionally, the two populations sorted from the healthy donors’ PBMCs, aforementioned, were cultured separately with MCF-7 spheroids (1:1). This co-culture was also maintained for 4 days, after which the spheroids were collected, stained and the viability of MCF-7 cells in each condition was assessed as above described.

### Statistical analysis

Statistical analysis was performed in GraphPad Prism v6 and SPSS v25 (IBM). Comparison between samples was performed by nonparametric Mann-Whitney test. To assess the biomarker performance, ROC curves were performed to assign a threshold to divide NACT-responders from non-responders. This cut-off point corresponded to the maximum of sensitivity and specificity. Univariate and multivariate logistic regressions were conducted taking into account both cohorts in the same analysis. Progression-free survival was analyzed by a Kaplan Meier curve with a logrank test and hazard ratio analysis. The probability model was elaborated in R (12). Statistical significance was considered for *p*<0.05.

## Results

### Clinical validation of HLA-DR level in cytotoxic T cells (CTLs) as a predictive biomarker of response to NACT

The characteristics of the patients selected to performed NACT (1^st^ cohort (9) and 2^nd^ cohort, flowchart in Fig.1) are described in Table 1.

We have previously observed, in the pilot study with 30 breast cancer (BC) patients selected for NACT, that HLA-DR level in CTLs was a putative predictive biomarker for the response to this treatment (9). Following the REMARK criteria (10), we have enrolled 31 patients in a 2^nd^ independent cohort to validate this biomarker (Fig.2). Response to NACT was defined as before (9). As in the first cohort, in this new cohort, although the percentage of total CTLs in the biopsies was identical between NACT-responders and non-responders (Fig.2A), patients with response to NACT had higher expression of HLA-DR in intratumor CTLs when compared to NACT non-responders (*p*<0.0001, Fig.2B). Then, following the REMARK criteria, we performed a series of statistical analyses, namely ROC curve, univariate and multivariate regressions.

**Fig.2.**
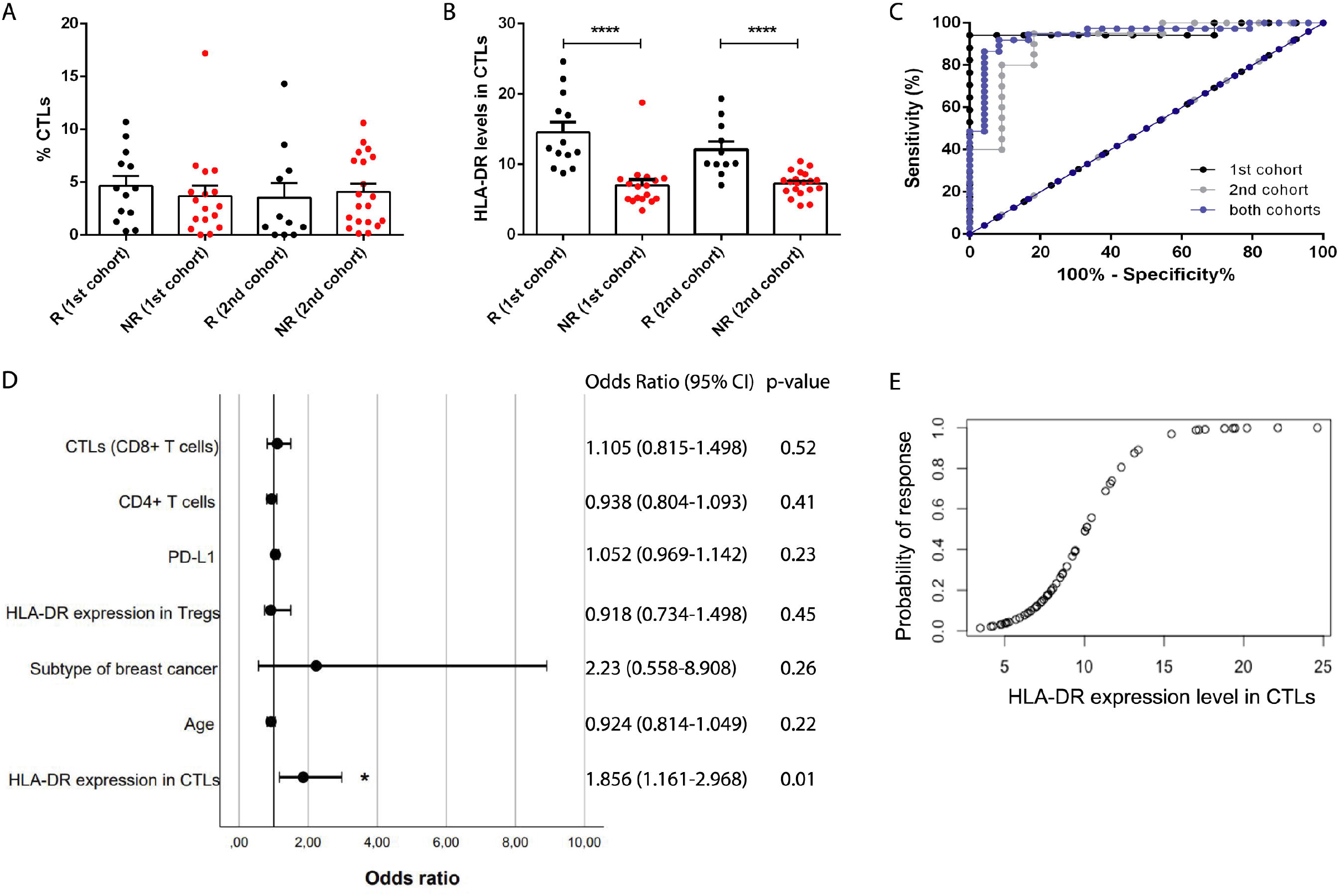
HLA-DR level in cytotoxic T cells is a clinically validated predictive biomarker of breast cancer patients’ response to NACT. **(A)** Percentage of infiltrating cytotoxic T cells (CTLs) in NACT-responders (black dots, 1^st^ cohort n=13 and 2^nd^ cohort n=11) and non-responders (red dots, 1^st^ cohort n=17 and 2^nd^ cohort n=20) breast cancer patients. **(B)** HLA-DR expression level in CTLs in the same patients as in (A). **(C)** ROC curve analysis of HLA-DR level in CTLs for the 1^st^ cohort (black lines), 2^nd^ cohort (grey lines) and merged cohorts (blue lines). **(D)** Forest plot of the multivariate analysis performed with both cohorts, including the odds ratio (with 95% confidence interval) and the *p*-values. **(E)** Predictive probability model of response to NACT according to the HLA-DR level in CTLs (merged cohorts). \*\*\*\**p*<0.0001.

ROC curve was executed to determine the cut-off point of HLA-DR level in CTLs to segregate NACT-responders from non-responders (Fig.2C). In the ROC curve of the first cohort (9), this value was 8.94, with an area under the curve (AUC) of 0.96, 94.12% sensitivity and 100% specificity (Fig.2C). Remarkably, in the ROC curve of the 2^nd^ cohort, we obtained similar parameters, namely a threshold value of 8.63, an AUC of 0.91, 80% sensitivity and 90.91% specificity (Fig.2C), hence validating our biomarker as a predictor of response to NACT.

Then, to pursue with the statistical analyses we joined the data regarding both cohorts. The resultant ROC curve had a cut-off value of 9.32, an AUC of 0.95, 91.89% sensitivity and 91.67% specificity (Fig.2C), which were very strong parameters, thus corroborating the robustness of HLA-DR level in CTLs to discriminate NACT-responders from non-responders.

By univariate analysis, we have observed that the biomarker HLA-DR level in CTLs was the only significant variable in predicting response to NACT (*p*<0.0001, OR=1.965 (95% CI 1.35-2.86), Table S1), in a multitude of clinical data and other immunological features assessed. While by multivariate analysis (represented by the forest plot in Fig.2D), we observed that this biomarker can predict BC patients’ response to NACT (*p*=0.01, OR=1.856 (95% CI 1.161-2.968)) even when the patients’ age, the BC subtype and other immunological parameters of the tumor, such as HLA-DR expression level in Tregs, PD-L1 expression in tumor cells, percentage of infiltrating CD4+ and CD8+ T cells are taken into account.

Since we have previously demonstrated that there was a correlation between the HLA-DR level in systemic and intratumor CTLs (9), here we used another cohort of BC patients (41 non-matched blood samples) and observed that NACT-responders present higher levels of HLA-DR in circulating CTLs than non-responders (*p*=0.03, Fig.S1), attesting the idea that this biomarker is reflected in circulation. Nonetheless, the ROC curve parameters, in this case, were not so strong as for the intratumor HLA-DR levels in CTLs (cut-off value of 15.72, AUC of 0.7, 85% of sensitivity and 52.38% of specificity).

Therefore, anticipating a potential clinical implementation of this biomarker to determine which patients will benefit from NACT and which should be promptly directed to surgery (if possible) or to alternative treatments, we have developed a predictive probability model, based on the analysis of this biomarker in BC patients’ biopsies (Fig.2E), to guide wiser therapeutic decisions. Indeed, following the equation

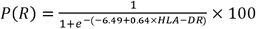

where P(R) is the probability of response and HLA-DR is the level of this marker in CTLs, it is possible to assess, in advance, the likelihood of a BC patient to respond to NACT, considering the biomarker here described.

### HLA-DR level in cytotoxic T cells (CTLs) is also a prognostic factor for breast cancer

Since the expression level of HLA-DR in CTLs is related with the activation status of these cells, we hypothesized that besides being a predictive biomarker of NACT response, it could also be useful as a prognostic factor, related with patients’ clinical outcome. As such, we have enrolled both non-NACT and NACT patients and performed a follow-up of 34 months. Dividing patients with HLA-DR^low^ CTLs and with HLA-DR^high^ CTLs (assessed in the biopsies or surgical specimens) according to the threshold value calculated in the ROC curve (considering both cohorts, Fig.2C), we observed a significant relationship between the level of HLA-DR in CTLs and the progression-free survival (PFS) curve (*p*=0.02, Fig.3). Namely, patients with HLA-DR^low^ CTLs have a lower PFS (Hazard Ratio (HR)=4.98 (95% CI 1.54-10.31)) when compared to patients with HLA-DR^high^ CTLs (HR=0.2 (95% CI 0.1-0.65), Fig. 3) Thus, we propose that patients with HLA-DR^low^ CTLs progress or relapse sooner than patients with HLA-DR^high^ CTLs, regardless of the treatment.

**Fig.3.**
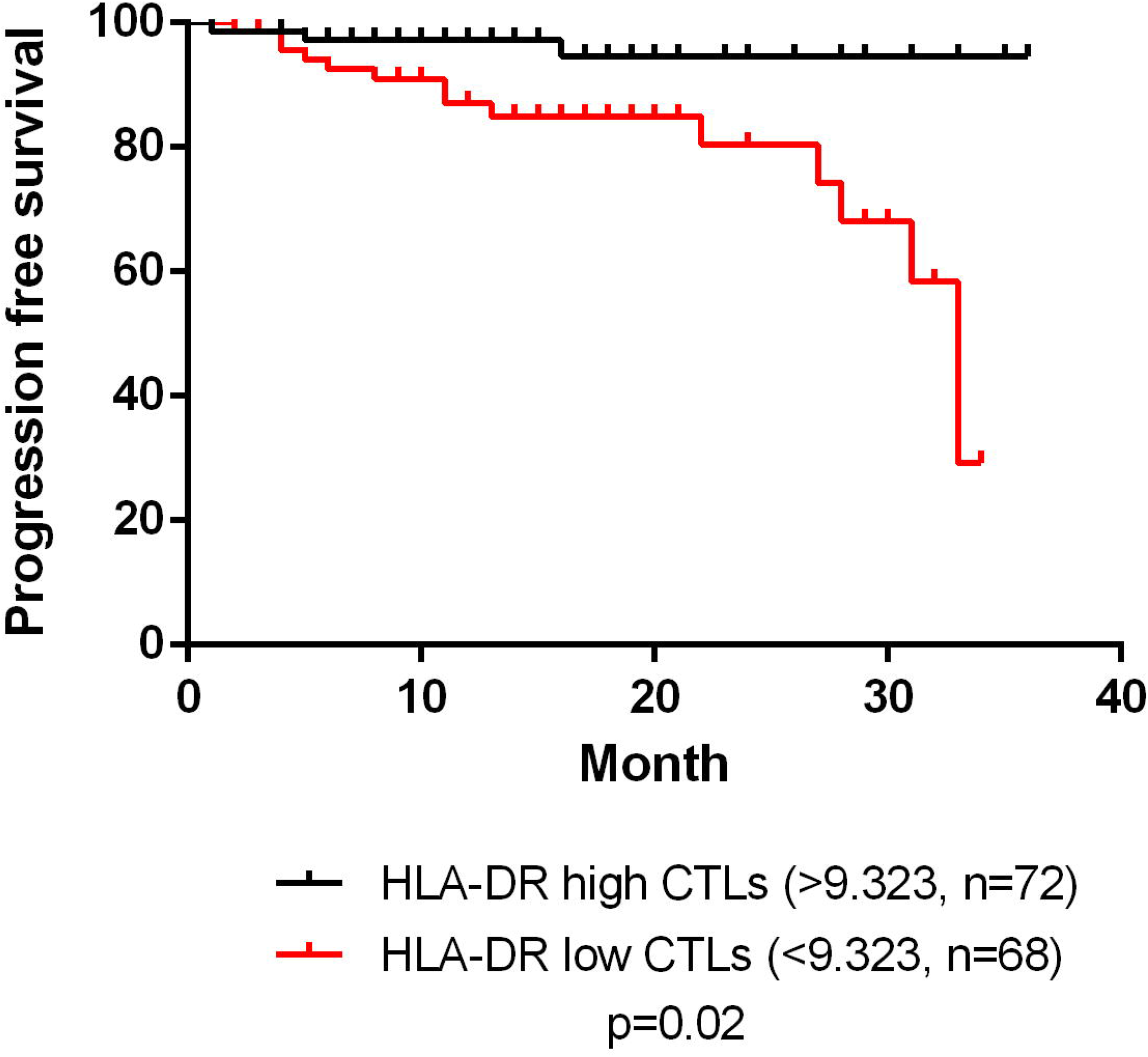
HLA-DR level in cytotoxic T cells has prognostic value. Progression-free survival of breast cancer patients divided in HLA-DR high in cytotoxic T cells (CTLs) (black line, n=72) and HLA-DR low in CTLs (red line, n=68); this segregation was performed according to the threshold value given by the ROC curve with data from both cohorts.

A similar analysis was performed for the overall survival (OS), but no differences were found in this case (data not shown), probably because for the OS a longer follow-up should be performed.

### HLA-DR+CTLs express high levels of activation and proliferation markers, release effector molecules and are cytotoxic against tumor cells

To shed some light on the reason why higher levels of HLA-DR in the tumor-infiltrating CTLs are required for the achievement of a good response to NACT, and eventually a better prognosis, we tried to better characterized the profile of HLA-DR+ CTLs in comparison with HLA-DRnegative CTLs (Fig.4A). By flow cytometry analysis, we observed that, when comparing to HLA-DRnegative CTLs, HLA-DR+ CTLs were more activated (CD69, *p*<0.0001) and proliferative cells (Ki67, *p*<0.0001) that produced more cytotoxicity-related molecules (IFN-γ and Granzyme B, *p*<0.0001), showed tissue-residency (CD103 and CD39, *p*<0.0001) and memory (CD45RO, *p*<0.0001) features, with similar levels of PD-1 and increased expression of Tim3 (p<0.01).

**Fig.4.**
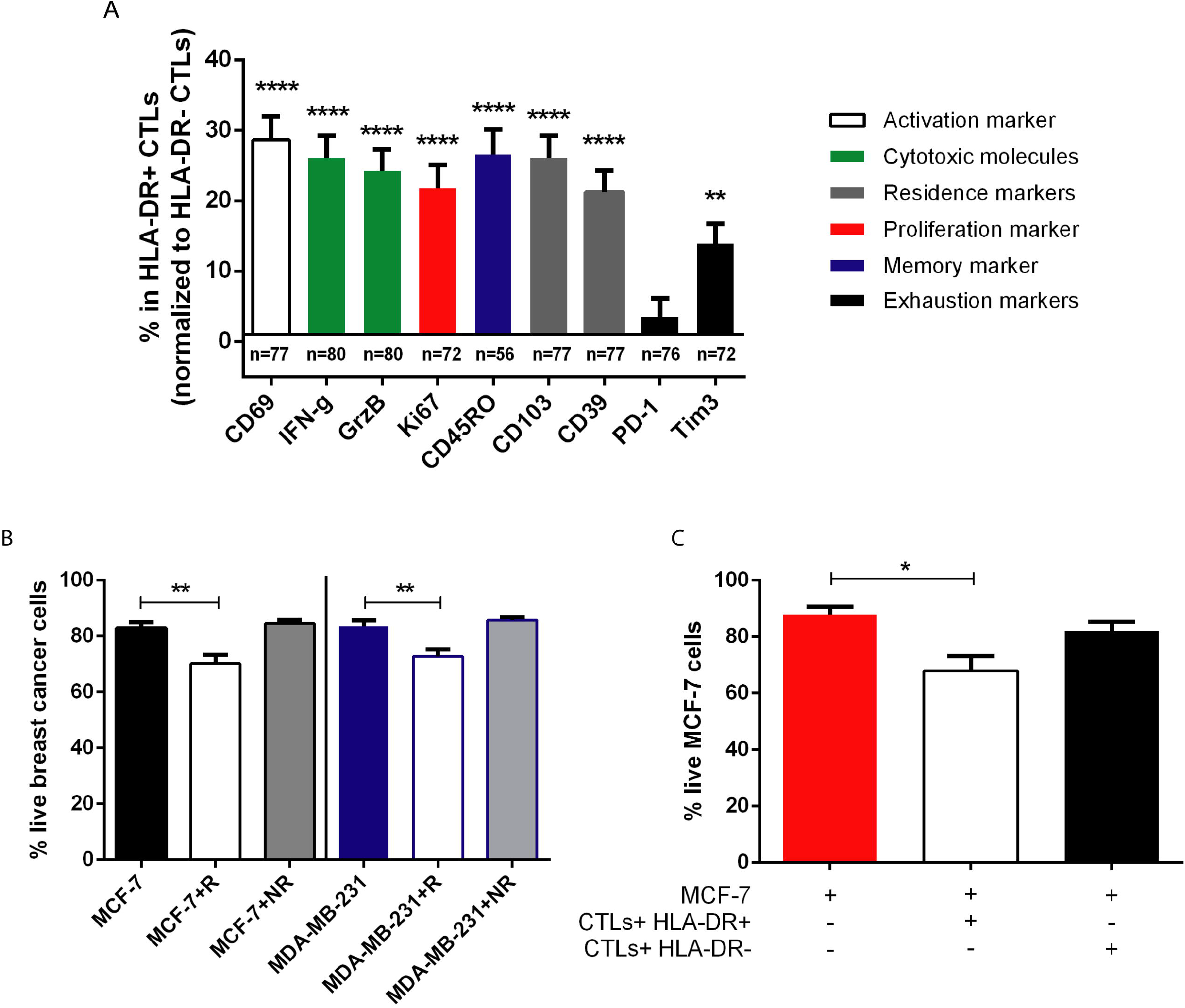
HLA-DR+ CTLs have enhanced anti-tumor properties in comparison to HLA-DRnegative CTLs. **(A)** Percentage of the activation marker CD69, the cytotoxic molecules IFN-γ and Granzyme B (GrzB), the proliferation marker Ki67, the memory marker CD45RO, the tissue-resident markers CD103 and CD39, and the exhaustion markers PD-1 and Tim3 in tumor infiltrating HLA-DR+ CTLs, normalized to HLA-DR-CTLs (number of biopsies used for this characterization is described in the graph). **(B)** Percentage of live MCF-7 cells in monoculture (MCF-7, black bar, n=10), in co-culture with NACT-responders’ PBMCs (MCF-7+R, white bar, n=10), and with NACT nonresponders’ PBMCs (MCF-7+NR, grey bar, n=11); percentage of live MDA-MB-231 cells in monoculture (MDA-MB-231, blue bar, n=11), in co-culture with NACT-responders’ PBMCs (MDA-MB-231+R, white bar, n=10), and with NACT nonresponders’ PBMCs (MDA-MB-231+NR, grey bar, n=12). **(C)** Viability of MCF-7 cells either in monoculture (red bar, n=5) or in co-culture with sorted HLA-DR+ CTLs (white bar, n=8) and HLA-DR negative CTLs (black bar, n=8). Data are represented as mean ± SEM, \**p*<0.05, \*\**p*<0.01, \*\*\*\**p*<0.0001.

Then, and since we demonstrated that in blood from NACT-responders, CTLs also exhibit higher levels of HLA-DR than CTLs from NACT non-responders ((9), Fig.S1) we used patient-derived peripheral blood mononuclear cells (PBMCs) isolated from the blood to perform *ex vivo* assays.

With the 3D *in vitro* platform we have developed (11), we added to two different BC cell lines (MCF-7 and MDA-MB-231) PBMCs from NACT-responders and non-responders, and assessed the viability of the tumor cells when in contact with patient-derived PBMCs (Fig.4B). Interestingly, we noticed that the addition of NACT-responders’ PBMCs (that are enriched in HLA-DR^high^ CTLs (Fig.S1) to the BC 3D-structure, reduced their viability after 4 days in co-culture (*p*=0.04 and *p*=0.06, Fig.4B); whereas the addition of non-responders’ PBMCs (that lack HLA-DR^high^ CTLs) showed no effect on the viability of the BC cells (Fig.4B), recapitulating somehow the clinical observations (Fig.2 and Fig.S1).

Taking into consideration that HLA-DR+ CTLs are more activated, proliferative and produce more cytotoxicity-related molecules (Fig.4A), we hypothesized that HLA-DR+ CTLs indeed exhibit anti-tumor properties and are cytotoxic against tumor cells, unlike the HLA-DRnegative CTLs. To confirm this, we used sorted HLA-DR+ and HLA-DRnegative CTLs, isolated from the same healthy individuals, after stimulation of their PBMCs, and cultured them separately with 3D structures of MCF-7 BC cells (Fig.4C). After 4 days in co-culture, we observed that the BC viability was reduced only in the presence of HLA-DR+ CTLs (*p*=0.02, Fig.4C), emphasizing that these cells, and not HLA-DRnegative CTLs, are cytotoxic against tumor cells.

## Discussion

Biomarkers that can effectively predict breast cancer (BC) patients’ response to NACT have been widely searched for, due to the unmet clinical need to determine which patients would not benefit from this treatment, in order to promptly direct them to more targeted approaches, avoiding the chemotherapy-related toxicity and the misuse of resources.

Tumor-infiltrating lymphocytes (TILs) have been studied as predictive biomarkers of the response to neoadjuvant chemotherapy (NACT) (13–16). However, TILs assessment is still not used in the clinical routine, despite an international effort to standardize the evaluation of this biomarker (17). The main disadvantages of the employment of TILs is the fact that they are presented as a single population, though they comprise immune cells with opposite roles (pro and anti-tumor T cells). Indeed, TILs represent a heterogeneous population, including the immunosuppressive regulatory T cells. Additionally, tumors can escape the immune system through, for instance, the release of anti-inflammatory cytokines (such as IL-10 or TGF-β), or the activation of immune checkpoints, namely the PD-1/PD-L1 axis, that will prevent the activation of CTLs and, consequently, lowering their cytotoxic function. Only when these escape mechanisms are not yet engaged, CTLs can become activated by recognizing tumor antigens and produce effector molecules, such as Perforin and Granzyme B, that leads to tumor cell elimination. Moreover, TILs have been suggested mainly for TNBC patients (excluding the majority of BC cases), its analysis is based on immunohistochemistry (IHC) of a single tumor plane, excluding the whole 3D conformation and, likewise, TIL scoring is more subjective.

To overcome the limitations of TILs, we previously proposed (9) and validate in the current study, a new biomarker – HLA-DR level in CTLs, which discriminates accurately NACT-responders from non-responders and better reflects the overall immune status of the tumor microenvironment. Actually, HLA-DR+ CTLs found in the biopsies represents a population of activated CTLs, as HLA-DR *per se* is an activation marker of T lymphocytes (18), but also expresses more of other activation markers (e.g. CD69) when compared with HLA-DRnegative CTLs. This subset also has higher proliferative capacity, more tissue-residency and memory markers (which have been highlighted as an interesting anti-tumor phenotype (19)), a similar level of exhaustion and, notably, increased cytotoxic properties against tumor cells (as observed in the 3D co-cultures), when compared to other CTLs present in the tumor microenvironment.

Thus, we verified, in two independent cohorts, that HLA-DR level in CTLs is, statistically, by the ROC curve parameters and the univariate and multivariate analysis of the merged data, a powerful predictive factor to segregate BC patients that will or will not respond to NACT. Additionally, this biomarker is independent of the BC subtype and other clinical parameters, such as the patients’ age, reflecting the capacity to be used in all BC patients that would be, based on the current criteria, selected to NACT.

Here, we proposed the assessment of HLA-DR expression level in CTLs by flow cytometry, and validated its determination in the following conditions: biopsies should be collected in Transfix (to preserve cellular antigens), processed and stained using a combination of two monoclonal mouse anti-human antibodies, specifically anti-CD8-PE (clone HIT8a) and anti-HLA-DR-APC (clone L243) from Biolegend in a 1:50 concentration. The flow cytometry analysis should be performed by gating HLA-DR within the CD8+ population (CTLs) and the value HLA-DR level in CTLs was obtained by determining the ratio between the median fluorescence intensity (MFI) of HLA-DR+ and the MFI of HLA-DRnegative population, as we previously reported (9). The employment of flow cytometry could be an advantage because we cannot distinguish intratumor CTLs from stromal CTLs, as it is possible with IHC, hence this technique is more representative of the whole biopsy than IHC, where only a slice of the 3D structure of the biopsy is used. Actually, several reports claim that stromal TILs are more important than the tumor-infiltrating ones to predict response to NACT (reviewed in (20)); while others report the opposite (21). With flow cytometry, we can quantify both simultaneously. Nevertheless, flow cytometry is not as widely available in Pathology departments of hospitals and clinics, as IHC is, which might represent a limitation.

Another advantage of the proposed biomarker is its systemic reflection, as NACT-responders also have higher expression of HLA-DR in blood CTLs. Nonetheless, the parameters of the ROC curve using blood samples were not as noticeable as for the biopsies, hence the prediction of NACT response should be assessed in the tissue biopsies. Yet, blood analysis could be useful as a complement and/or can be performed, for instance, to monitor NACT efficiency throughout the treatment.

Envisioning the application of this biomarker in a clinical setting, we have developed a probability model, which can be used by clinicians in the therapeutic decision process. Based on this model, we suggest segregating patients into 3 groups – high probability of response to NACT (>60% that corresponds to an HLA-DR level higher than 10.8), intermediate probability of response (20-59% that corresponds to an HLA-DR between 8 and 10.8) and low probability of response (<20% that corresponds to an HLA-DR level lower than 8). This stratification should help to guarantee that patients with high probability to respond to NACT will be submitted to this treatment; while patients with low probability to respond to NACT will be promptly directed to surgery/alternative therapies, whenever possible. Of course for the cases of patients with intermediate probability of response, the therapeutic decision would be more challenging and clinicians should analyze it case-by-case and contemplate different aspects, such as if another therapeutic option is available or if NACT is still the best option. Noteworthy, the equation presented was extrapolated from the results of these studies, using both cohorts merged, so it is only valid for HLA-DR level in CTLs ranging from 4.13 to 24.63, determined as we did.

Recently, we have developed a 3D platform composed of BC cell lines and PBMCs (11), anticipating the use of this system as a patient-customized drug screening assay. Here, we took advantage of this 3D system to demonstrate *in vitro* that only PBMCs from NACT-responders have the ability to reduce the viability of tumor cells. Given the fact that HLA-DR expression is boosted in circulating CTLs from NACT-responder patients, we further showed that it is this subset, and not other CTLs, that in fact are responsible for the anti-tumor properties of NACT-responders’ PBMCs. In line with the idea that an immune status favoring cytotoxicity against tumors will determine the success of chemotherapy, by allowing the control of chemo-resistant cancer cells, these results contribute to explain the requirement of HLA-DR^high^ CTLs for NACT efficacy and to corroborate clinical observations.

Besides being a predictive factor, HLA-DR level in CTLs is also associated with the progression-free survival curve considering 34 months of follow-up of all the patients whose biopsies/surgical specimens we have analyzed, independently of the treatment performed. This observation suggests that besides the predictive value of this biomarker for the response to a specific treatment, it has also a promising prognostic value that could be useful to determine, in advance, BC patients’ outcome.

Furthermore, this biomarker also has the potential to be used in clinical trial design to evaluate highly expensive and technically demanding immunotherapies, particularly in those BC patients who might truly need/take advantage of these novel treatments. In the future, it may be also adapted as a predictive/prognostic factor for other types of cancer.

## Acknowledgments

We would like to thank all the breast cancer patients that agreed to participate in this study. We also appreciate all the help from the Radiologists, Pathologists, Oncologists, Surgeons and Nurses from Hospital CUF Descobertas, Hospital CUF Tejo, Hospital de Vila Franca de Xira, Hospital Prof. Doutor Fernando Fonseca and Hospital Santa Maria in gathering the samples. We also acknowledge the support of the Flow Cytometry Facility at CEDOC.

## Author contributions

DPS conducted all the experiments, analyzed and interpreted the data, performed the statistical analysis, assembled all the figures and wrote the manuscript. SAL and AA helped with the statistical analysis. RS helped with flow cytometry experiments and in obtaining clinical data. PB, BA, IP, ZS, IN and SB helped in the obtainment of patients’ samples. AJ contributed to scientific discussion. MGC designed and supervised the study, interpreted the data and wrote the manuscript. All the authors approved the final manuscript.

## References

1. Bray F, Ferlay J, Soerjomataram I, Siegel RL, Torre LA, Jemal A. Global cancer statistics 2018: GLOBOCAN estimates of incidence and mortality worldwide for 36 cancers in 185 countries. CA: A Cancer Journal for Clinicians. 2018 Nov;68(6):394–424.

2. Waks AG, Winer EP. Breast Cancer Treatment: A Review. JAMA. 2019 Jan 22;321(3):288.

3. Thompson AM, Moulder-Thompson SL. Neoadjuvant treatment of breast cancer. Annals of Oncology. 2012 Sep;23:x231–6.

4. Cortazar P, Zhang L, Untch M, Mehta K, Costantino JP, Wolmark N, et al. Pathological complete response and long-term clinical benefit in breast cancer: the CTNeoBC pooled analysis. The Lancet. 2014 Jul;384(9938):164–72.

5. DeMichele A, Yee D, Esserman L. Mechanisms of Resistance to Neoadjuvant Chemotherapy in Breast Cancer. Phimister EG, editor. N Engl J Med. 2017 Dec 7;377(23):2287–9.

6. Fuchs Y, Steller H. Live to die another way: modes of programmed cell death and the signals emanating from dying cells. Nat Rev Mol Cell Biol. 2015 Jun;16(6):329–44.

7. Krysko O, Løve Aaes T, Bachert C, Vandenabeele P, Krysko DV. Many faces of DAMPs in cancer therapy. Cell Death Dis. 2013 May;4(5):e631–e631.

8. O’Donnell JS, Teng MWL, Smyth MJ. Cancer immunoediting and resistance to T cell-based immunotherapy. Nat Rev Clin Oncol. 2019;16(3):151–67.

9. Saraiva DP, Jacinto A, Borralho P, Braga S, Cabral MG. HLA-DR in Cytotoxic T Lymphocytes Predicts Breast Cancer Patients’ Response to Neoadjuvant Chemotherapy. Front Immunol. 2018 Nov 13;9:2605.

10. McShane LM, Sauerbrei W, Taube SE, Gion M, Clark GM. REporting recommendations for tumour MARKer prognostic studies (REMARK). Br J Cancer. 2005 Aug;93(4):387–91.

11. Saraiva DP, Matias AT, Braga S, Jacinto A, Cabral MG. Establishment of a 3D Co-culture With MDA-MB-231 Breast Cancer Cell Line and Patient-Derived Immune Cells for Application in the Development of Immunotherapies. Front Oncol. 2020 Aug 27;10:1543.

12. R: A language and environment for statistical computing. R Foundation for Statistical Computing [Internet]. 2020; Available from: https://www.R-project.org/

13. Russo L, Maltese A, Betancourt L, Romero G, Cialoni D, De la Fuente L, et al. Locally advanced breast cancer: Tumor-infiltrating lymphocytes as a predictive factor of response to neoadjuvant chemotherapy. European Journal of Surgical Oncology. 2019 Jun;45(6):963–8.

14. Denkert C, von Minckwitz G, Brase JC, Sinn BV, Gade S, Kronenwett R, et al. Tumor-Infiltrating Lymphocytes and Response to Neoadjuvant Chemotherapy With or Without Carboplatin in Human Epidermal Growth Factor Receptor 2– Positive and Triple-Negative Primary Breast Cancers. JCO. 2015 Mar 20;33(9):983–91.

15. Denkert C, Loibl S, Noske A, Roller M, Müller BM, Komor M, et al. Tumor-associated lymphocytes as an independent predictor of response to neoadjuvant chemotherapy in breast cancer. J Clin Oncol. 2010 Jan 1;28(1):105–13.

16. Loibl S, Untch M, Burchardi N, Huober J, Sinn BV, Blohmer J-U, et al. A randomised phase II study investigating durvalumab in addition to an anthracycline taxane-based neoadjuvant therapy in early triple-negative breast cancer: clinical results and biomarker analysis of GeparNuevo study. Ann Oncol. 2019 01;30(8):1279–88.

17. Salgado R, Denkert C, Demaria S, Sirtaine N, Klauschen F, Pruneri G, et al. The evaluation of tumor-infiltrating lymphocytes (TILs) in breast cancer: recommendations by an International TILs Working Group 2014. Annals of Oncology. 2015 Feb;26(2):259–71.

18. Rea IM, McNerlan SE, Alexander HD. CD69, CD25, and HLA-DR activation antigen expression on CD3+ lymphocytes and relationship to serum TNF-α, IFN-γ, and sIL-2R levels in aging. Experimental Gerontology. 1999 Jan;34(1):79–93.

19. Duhen T, Duhen R, Montler R, Moses J, Moudgil T, de Miranda NF, et al. Coexpression of CD39 and CD103 identifies tumor-reactive CD8 T cells in human solid tumors. Nat Commun. 2018 Dec;9(1):2724.

20. Pruneri G, Vingiani A, Denkert C. Tumor infiltrating lymphocytes in early breast cancer. The Breast. 2018 Feb;37:207–14.

21. Khoury T, Nagrale V, Opyrchal M, Peng X, Wang D, Yao S. Prognostic Significance of Stromal Versus Intratumoral Infiltrating Lymphocytes in Different Subtypes of Breast Cancer Treated With Cytotoxic Neoadjuvant Chemotherapy: Applied Immunohistochemistry & Molecular Morphology. 2017 Feb;1.

