## Supplementary material for "Determination of HLA-DR level in cytotoxic T lymphocytes: a new validated tool to predict breast cancer response to treatment": Table S1, Fig. S1

Table S1 - **Univariate** **analysis for the biomarker HLA-DR level in CTLs**.

HLA-DR expression level in CTLs and other immunological markers and clinical parameters were analyzed by univariate analysis. p-value, odds ratio and the confidence interval of the odds ratio are represented.

| Parameter | p-value | Odds Ratio | 95% Confidence Interval |
| --- | --- | --- | --- |
| HLA-DR expression level in CTLs | <0.0001 | 1.965 | 1.35-2.86 |
| Immune cells (CD45) | 0.985 | 1.000 | 0.965-1.036 |
| T cells (CD3) | 0.520 | 0.985 | 0.94-1.032 |
| B cells (CD19) | 0.919 | 0.995 | 0.904-1.095 |
| NK cells (CD161) | 0.122 | 0.852 | 0.696-1.044 |
| T helper cells (CD4) | 0.282 | 0.972 | 0.922-1.024 |
| CTLs (CD8) | 0.934 | 1.006 | 0.875-1.156 |
| Tregs (CD25^high^/CD127^low^) | 0.503 | 1.157 | 0.755-1.772 |
| M1 macrophages (CD11b+/CD163-/CD206-) | 0.719 | 0.628 | 0.05-7.944 |
| M2 macrophages (CD11b+/CD163+/CD206+) | 0.743 | 0.939 | 0.644-1.369 |
| Neutrophils (CD15) | 0.298 | 0.897 | 0.731-1.101 |
| HLA-DR expression level in Tregs | 0.724 | 0.981 | 0.885-1.089 |
| PD-L1 | 0.165 | 1.015 | 0.994-1.037 |
| IL-10 | 0.107 | 1.051 | 1-1.104 |
| Age | 0.254 | 0.975 | 0.933-1.018 |
| Body Mass Index | 0.69 | 0.977 | 0.869-1.097 |
| Tumor dimension | 0.167 | 1.024 | 0.99-1.06 |
| Node invasion status | 0.338 | 0.566 | 0.177-1.813 |
| Ki67 (%) | 0.271 | 1.012 | 0.991-1.033 |
| Grade | 0.474 | 0.563 | 0.116-2.718 |

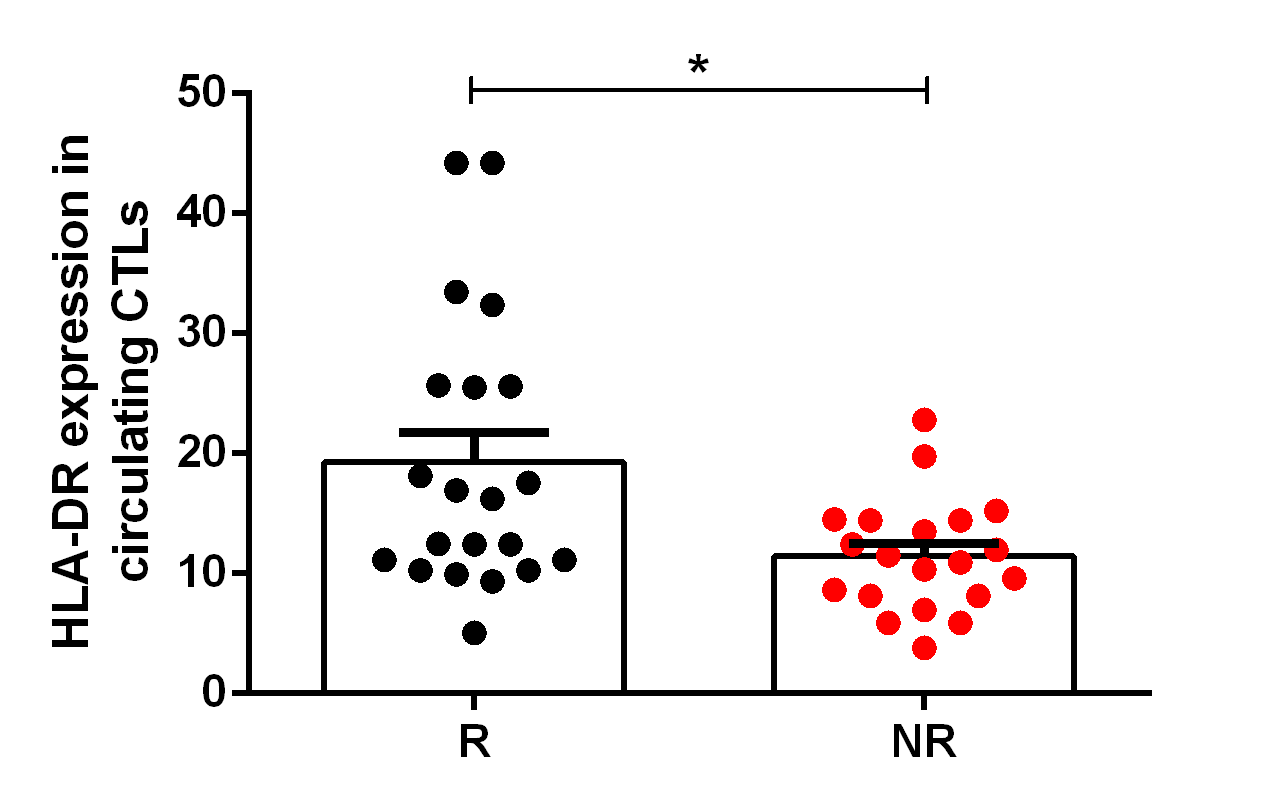

Fig.S1 – **HLA-DR expression level in systemic cytotoxic T cells segregate breast cancer patients according to their response to neoadjuvant chemotherapy.** HLA-DR expression level in circulating cytotoxic T lymphocytes (CTLs) of NACT-responders (R, black dots, n=21) and NACT non-responders (NR, red dots, n=20). *p<0.05.
